# Virus replication in the honey bee parasite, *Varroa destructor*

**DOI:** 10.1101/2023.07.16.549232

**Authors:** James E. Damayo, Rebecca C. McKee, Gabriele Buchmann, Amanda M. Norton, Alyson Ashe, Emily J. Remnant

## Abstract

Arthropod vectors such as mites and ticks introduce an alternative viral transmission route between their hosts. The ectoparasitic mite *Varroa destructor* is the leading threat to the health of Western honey bees (*Apis mellifera*) primarily through its action as a vector of viruses. However, it is unclear whether viruses transmitted by *V. destructor* actively infect and replicate in mites, which could facilitate increased transmission and select for more virulent strains. To better understand the role of *V. destructor* as a vector, we took advantage of differences between bee and mite antiviral RNA interference pathways to identify the host specificity of replicating viruses. We used small RNA sequencing of individual *V. destructor* mites to examine viral small interfering RNA (vsiRNA) profiles of Deformed wing virus genotypes (DWV-A and DWV-B), associated with colony declines, as well as nine other viruses present in our samples. We found active replication of six *V. destructor*-associated viruses, including a novel virus, Varroa destructor virus 9 (VDV-9), and replication of two honey bee associated viruses, including both DWV-A and -B genotypes, suggesting that mites are biological vectors for important bee pathogens. We show that the antiviral RNAi response can be used define the host range of viruses in host-parasite interactions, such as honey bees and their parasites, enabling a better understanding of the role of a vector in the evolution and spread of honey bee pathogens.

## INTRODUCTION

Many viruses are generalist pathogens with a broad host range, able to infect and replicate in multiple hosts (Rothenburg and Brennan, 2020; Woolhouse et al., 2001). The host range of a virus depends on a variety of factors at both cellular and ecological scales. At a cellular level, a multi-host virus must circumvent molecular barriers to facilitate successful infection across different species, including receptor attachment to enable cellular entry (Maginnis, 2018), co-opting host replication machinery, and evading or overcoming the host immune response (Woolhouse et al., 2001). At ecological scales, the geographic distribution and degree of shared ecological contact between alternative host species can determine the nature and frequency of exposures to multi-host viruses (Parrish et al., 2008). Disease transmission dynamics between multi-host viruses are often complex (Webster et al., 2017). Prolonged exposure to a virus via an infected donor host can lead to greater opportunities for spillover and host switching into recipient hosts (Parrish et al., 2008). RNA viruses are particularly adept at switching hosts, due to high mutation and recombination rates driving rapid adaptations to enable replication in multiple organisms (Holmes, 2009b).

The intimate association between a parasite and host provides ample opportunity for reciprocal exchange of microbes, such as viruses. Cross-species transmission of viruses is exemplified in arthropod vectors such as mosquitoes and ticks, although active viral replication and infection may not always occur in both the vector and host. Some vectors are mechanical, transferring viral particles between hosts via contact without playing an active role in the virus replication cycle, such as the biting arthropods that carry myxoma virus between rabbits (Kerr et al., 2015). Alternatively, biological vectors like *Aedes* mosquitoes play an essential part of the infectious cycle of Dengue and Yellow fever viruses, where viral replication or maturation occurs within the vector, facilitating infectivity in human hosts (Mellor, 2000). Biological vectors may therefore require enhanced mechanisms to withstand viral infections and maintain fitness while harbouring high viral loads as a consequence of viral replication (Oliveira et al., 2020).

Western honey bees (*Apis mellifera*) are of critical importance to the agricultural industry as pollinators for a diverse range of horticultural crops (Calderone, 2012). Within the last few decades however, the health of honey bees has declined across most of the globe, exacerbated by the spread of exotic parasites and pathogens that has been facilitated by the international honey bee trade (Neumann and Carreck, 2010). In particular, movement of *A. mellifera* into Asia exposed Western honey bees to mites of the genus *Varroa,* obligate ectoparasites originally restricted to the Eastern hive bee, *Apis cerana* (Oldroyd, 1999). In the decades following, two mite species, *V. destructor* and *V. jacobsoni,* switched hosts from *A. cerana* to *A. mellifera,* with *V. destructor* in particular rapidly dispersing across the globe, driving widespread colony losses (Anderson and Trueman, 2000; Roberts et al., 2015; Rosenkranz et al., 2010).

Mites physiologically affect honey bees by feeding on fat body tissue of juvenile and adult bees, depleting protein, carbohydrate and water contents (Bowen-Walker and Gunn, 2001; Ramsey et al., 2019). Additionally, *V. destructor* is an efficient vector of viruses which it can acquire during feeding on infected honey bee hosts, increasing dissemination and dispersal of these viruses within and between colonies (Bowen-Walker et al., 1999; Ribière et al., 2008; Traynor et al., 2020). *Varroa* mites have drastically altered the honey bee viral landscape, as demonstrated during the spread of *V. destructor* throughout New Zealand, where dynamic shifts occurred in the prevalence and load of multiple common honey bee viruses in the years following *V. destructor* introduction (Mondet et al., 2014). Initial increases in Kashmir bee virus (KBV), Sacbrood virus (SBV) and Black queen cell virus (BQCV) were eventually superseded by Deformed wing virus (DWV), which now remains the predominant virus in New Zealand colonies (Lester et al., 2022). Similarly, in Hawaii, the introduction of *V. destructor* led to increased prevalence of DWV in managed colonies from ∼10% to 100% and an overall reduction in DWV genetic diversity, resulting in the selection of a single dominant DWV genotype (Martin et al., 2012). The association of DWV with *V. destructor* infestation is frequently implicated as a major driver of colony declines (Di Prisco et al., 2016; Wilfert et al., 2016), due to increased abundance of mites leading to higher DWV loads (Norton et al., 2021), which in turn is linked to weaker colonies (Barroso-Arevalo et al., 2019; Highfield et al., 2009).

DWV is an icosahedral, positive-sense, single-stranded RNA virus belonging to the *Iflaviridae* family, and exists as a multi-strain viral complex (Martin and Brettell, 2019). Two DWV genetic variants, DWV-A and DWV-B, are closely related with 85% nucleotide identity, and are both frequently identified at high abundance in *V. destructor-*infested colonies. An additional two genotypes have been characterised that are more distantly related; DWV-C (Mordecai et al., 2015) and DWV-D (’Egypt bee virus’; de Miranda et al., 2022); these variants are more restricted in their distribution or historical occurrence. Although DWV-A and -B genotypes are both widespread, they vary in their geographic distribution and their global prevalence has changed over time (Hasegawa et al., 2023). Historically, DWV-A was the most prevalent genotype, however over time DWV-B has increased and it is predicted that it will eventually surpass DWV-A (Kevill et al., 2021; Paxton et al., 2022; Ryabov et al., 2017). Frequent coinfections have led to multiple instances of recombination between DWV-A and -B genomes, resulting in the presence of stable recombinant genotypes (Cornman, 2017; Ryabov et al., 2014). The different genotypes and recombinants differ in characteristics such as viral accumulation, competition, replication rate and virulence (Norton et al., 2020; Paxton et al., 2022).

The mechanisms by which *Varroa* leads to increased viral prevalence and load are multifaceted. In the absence of *Varroa,* virus transmission between bees normally occurs via the faecal-oral route, leading to covert infections that rarely present with disease symptoms (Ribière et al., 2008). *Varroa* parasitism introduces an alternate, vector-mediated transmission route, facilitating direct injection of viral particles into the body, leading to systemic, overt infections that circumvent innate defences provided by the gut and cuticle (Evans and Spivak, 2010). Increased viral loads and disease symptoms could therefore directly result from passive, mechanical vectoring of viral particles via mite mouthparts or gut contents as mites sequentially feed on new hosts (Piou et al., 2022). However, it is becoming increasingly evident that *V. destructor* is also a biological vector for DWV, with strong evidence of active replication of DWV-B and recombinant DWV genotypes within *Varroa* tissues (Gisder and Genersch, 2021; Gusachenko et al., 2020). Replication of DWV within *Varroa* changes the evolutionary dynamics of virus transmission, leading to higher viral loads in mites, larger doses inoculated into bees, and selection for genotypes that can replicate in both vector and host, with potentially higher virulence in honey bees (Eliash et al., 2022). There remains some contention regarding whether all DWV genotypes, in particular DWV-A, are able to replicate in mites (Gisder and Genersch, 2021), with experimental evidence suggesting that DWV-A is transmitted by mites in a non-propagative manner (Posada-Florez et al., 2019).

In addition to DWV, *V. destructor* carries a diverse virome that may contain other honey bee-infecting viruses, along with a suite of *Varroa-*infecting viruses. Like most eukaryotes, *V. destructor-*associated viruses are primarily single stranded RNA viruses (Holmes, 2009a). At least 11 positive sense RNA viruses (+ssRNA) have been identified infecting *V. destructor* (de Miranda et al., 2015; Herrero et al., 2019; Levin et al., 2016; Levin et al., 2019; Li et al., 2023); as well as three negative sense RNA viruses (-ssRNA; Levin et al., 2019; Remnant et al., 2017), and three DNA viruses (Cornman et al., 2010; Gauthier et al., 2015; Kraberger et al., 2018). Some viruses such as Apis Rhabdovirus-1 (ARV-1), Apis Rhabdovirus-2 (ARV-2) and Varroa destructor virus-2 (VDV-2) are highly prevalent, found in virtually all *V. destructor* transcriptomes examined (Eliash et al., 2022; Lester et al., 2022).

The obligate association between *V. destructor* mites and their honey bee hosts makes it challenging to definitively identify which viruses are able to replicate in which species. Many studies attempt to identify RNA virus replication using strand-specific RT-PCR to detect the antigenome of replicating single-stranded RNA viruses, which produce double-stranded RNA virus intermediates during virus replication (Gisder et al., 2009; Ongus et al., 2004). While this method may be suitable in non-parasitic organisms, parasites such as *Varroa* ingest infected host tissue that can contain viral replication intermediates, which would provide false positives during strand-specific RT-PCR assays (Liu et al., 2011; Posada-Florez et al., 2019). More robust methods are therefore necessary to definitively identify which of the honey bee-infecting viruses can actively replicate in *V. destructor*. Fluorescent tagging of genetically engineered viral genomes (Gusachenko et al., 2020) and *in situ* hybridisation of negative-sense RNA replication intermediates (Gisder and Genersch, 2021) has been successfully used to assess DWV virus replication in mites, however this becomes increasingly unfeasible when assessing replication of multiple viruses, distinct viral variants, or novel and uncharacterised virus species.

One efficient and unbiased method that can identify all replicating viruses within a sample is small RNA sequencing and examination of the antiviral RNA interference (RNAi) response. RNAi is one of the main antiviral defence mechanisms in arthropods (Kingsolver et al., 2013). The antiviral RNAi pathway is triggered by the presence of dsRNA produced during viral replication, that is detected by the endonuclease Dicer-2, which then cleaves dsRNA into small interfering RNA (siRNA) 21-25 nt fragments (Jinek and Doudna, 2009). Cleaved siRNAs associate with the Argonaute protein AGO2, which delivers guide siRNAs to the RNA-Induced Silencing Complex (RISC). The guide siRNAs bind to complementary sequences in their equivalent viral genomes, leading to degradation of virus RNA and suppression of viral infection (Gammon and Mello, 2015).

Insects typically produce sense and antisense viral siRNAs (vsiRNAs) of 21-23 nt (Chejanovsky et al., 2014; Webster et al., 2015), while other metazoans exhibit more divergent antiviral RNAi responses, with variation in vsiRNA length (23-30nt) and polarity (eg. negative strand bias; Waldron et al., 2018). Mites including *V. destructor* carry all genes required for the antiviral RNAi pathway (Nganso et al., 2020), however vsiRNA size profiles are distinct from those observed in insects. For example, both *A. mellifera* and *V. destructor* exhibit an active and abundant vsiRNA profile in response to infection with the -ssRNA rhabdoviruses ARV-1 and -2, indicating active viral replication occurs within both species. However, *V. destructor* produces 24 nt vsiRNA fragments with a strong antisense bias, in contrast to the sense/antisense 22 nt vsiRNAs in *A. mellifera* (Remnant et al., 2017). The contrasting vsiRNA size profiles between *A. mellifera* and *V. destructor* provides a unique opportunity to distinguish active antiviral responses in mites from those in bees, and determine the viruses that actively replicate in *V. destructor* and are therefore genuine mite infections. Applying this method broadly to *Varroa* populations carrying a range of honey bee-infecting viruses and strains, such as different genotypes of DWV, allows us to assess which of these are subject to biological vectoring.

In this study, we analysed small RNA profiles of individual and pooled *V. destructor* mites from New Zealand, the Netherlands, South Africa, China and the USA to identify active vsiRNA profiles indicative of replicating viruses. In particular, we were interested in determining whether multiple DWV genotypes undergo active replication in mites, and whether other common honey bee-infecting viruses also produce active vsiRNA profiles in *V. destructor,* to better understand the status of mites as biological vectors of important honey bee pathogens.

## MATERIALS AND METHODS

### Sample collection and processing

*V. destructor* samples were collected from *A. mellifera* colonies in New Zealand (NZ), the Netherlands (NE) and South Africa (SA) during 2013-2018 (see Table S1 for sampling details). RNA was extracted from 15 individual mites and six pooled mite samples consisting of four mites, to generate 21 new samples for small RNA sequencing (Table S1). Total RNA was extracted using TRIzol reagent (ThermoFisher). Whole individual mites were extracted in 100 µl of Trizol and pooled mite sample volumes were scaled accordingly. Mites were homogenized in half the total TRIzol reagent with a sterile micropestle. The remaining TRIzol was added, followed by 1/5^th^ volume of chloroform. RNA was precipitated from the aqueous phase with ½ volume of isopropanol and 1 µl glycogen, and incubated at -20° C overnight. Following initial precipitation, RNA pellets were washed in 75% EtOH, and RNA resuspended in 6 µl H_2_O (single mites) or 25 µl H_2_O (pooled mites). RNA purity was confirmed with a Nanodrop spectrophotometer (ThermoFisher), and concentrations were quantified with the Qubit Bioanalyzer (ThermoFisher).

### Library preparation and sequencing

Small RNA (sRNA) libraries were generated using total RNA from single and pooled mites (100-600 ng) with the NEBNext Multiplex Small RNA Library Prep kit (NEB) following the manufacturers protocol. Small RNA libraries were purified with the Monarch PCR Cleanup Kit (NEB) prior to size selection to obtain the correct fragment sizes for small RNA analysis. Libraries were mixed with Novex 5X Hi-Density TBE Sample Loading Dye (Thermofisher), then loaded into a 6% TBE acrylamide gel with the Quick-Load pBR322 DNA-MspI Digest ladder and run for 65 minutes at 120 V. Gels were stained with SYBR Gold (Thermofisher) in 6% TBE buffer for 3 minutes, prior to size selection under a UV transilluminator. The region encompassing the small RNA size range of interest (15-35 nt, corresponding to the size ladder bands at 147 - 160 nt) was excised and purified by first passing through gel breaker tubes in DNA Gel elution buffer (NEB), incubated overnight at 4° C with shaking, then precipitated with 100% EtOH, 3M sodium acetate (pH 5.5) and 2 µl glycogen for 4 hours at -80° C. The pellets were resuspended in 11 µl of TE buffer, and the sRNA libraries were shipped to the Australian Genome Research Facility (AGRF) for sequencing with Illumina HiSeq or Illumina NovaSeq 6000 (100bp SE). Raw sequence reads generated from this study have been deposited at the NCBI Sequence Read Archive under the BioProject PRJNA986961 (Table S1).

### Small RNA composition analysis

In addition to the 21 new samples produced in this study, we obtained additional small RNA libraries of seven previously sequenced *V. destructor* samples from the NCBI Sequence Read Archive, two from our previous study of South African *V. destructor* (Remnant et al., 2017), three from China (Li et al., 2023) and two from the USA (Kumar et al., 2022; Table S1). We initially examined all eight sRNA libraries from Kumar et al (2022), however discarded data from six due to evidence of pseudoreplication (McKee; unpublished). Sequencing adaptors and low quality reads were removed with TrimGalore (Krueger, 2022) and read quality was evaluated with FastQC (Andrews, 2010). To identify the sRNA composition of each sample, we first filtered our sequences for ribosomal RNA, because the proportion of rRNA reads captured during small RNA library synthesis is highly variable between samples (McKee, unpublished). We then performed sequential bowtie2 (Langmead and Salzberg, 2012) alignments, using the ‘--very sensitive local’ parameters, changing -L to 15 to facilitate short read mapping. Reads were first aligned to the *V. destructor* genome (Vdes_3.0; Genbank accession GCA_002443255.1), then the *A. mellifera* genome (Amel_HAv3.1; GCA_003254395.2), and finally to a reference library containing virus genomes associated with *V. destructor*, adapted from Lester *et al*. 2022 (Table S2). For each consecutive alignment, unmapped reads were written to a separate fastq file which was used for subsequent alignments. To identify shared reads between *V. destructor* and *A. mellifera*, libraries were first aligned to the *V. destructor* genome and the aligned reads were written to a separate file, and then mapped to the *A. mellifera* genome, with any aligned reads corresponding to the number of reads shared between the two genomes. To ensure the number of shared reads was consistent, reciprocal alignments starting with the *A. mellifera* genome, followed the *V. destructor* genome were performed. Resulting SAM files were converted to BAM files with Samtools (Li et al., 2009). Geneious Prime 2022.2.2 (https://www.geneious.com/) was used to visualise the BAM file alignments.

### Consensus virus sequence assembly

To obtain strain-specific virus reference genomes for more accurate siRNA alignments, we attempted to generate consensus genome sequences of the viruses in individual samples to identify viral strain polymorphisms using two methods. First, we performed a *de novo* assembly of small RNA reads for each sample using Megahit (v 1.2.9; Li et al., 2015). Assembled contigs were examined for viral homology using a BLASTx search to a non-redundant viral protein database (NCBI; accessed Aug 2022). With this approach we identified short fragments (∼200-6000 nt) of a number of known *V. destructor* and *A. mellifera* viruses, indicating that *de novo* assembly of our small RNA reads was generally insufficient for full viral genome reconstruction. However, in one small RNA library from the Netherlands (NE-6), BLASTx revealed two contigs with ∼30% homology to Beihai Horseshoe crab virus 1 (YP_009333375.1), indicating the presence of a putatively novel virus.

As we were not able obtain sufficient strain information for all virus genomes from *de novo* assembly, we generated additional consensus sequences by performing a series of iterative bowtie2 alignments of the short reads to reference viral genomes. We first performed an initial alignment to our reference virus genome library, and generated consensus viral sequences for each sample in Geneious Prime using the ‘Generate Consensus Sequence’ function. We set the consensus caller threshold in Geneious Prime to 0% ‘Majority’, to incorporate any sample-specific variants by allowing fewest nucleotide ambiguities, and if there was no coverage, the reference sequence was called. Any within-sample nucleotide ambiguities were manually resolved by randomly selecting a representative alternate nucleotide according to the International Union of Pure and Applied Chemistry (IUPAC) nucleotide naming nomenclature, prior to subsequent bowtie2 alignments. For subsequent iterative alignments, the sRNA reads from each sample were aligned to their corresponding consensus viral reference sequences with bowtie2, and new consensus viral sequences were synthesised using the aforementioned methods. Consensus generation and re-alignment was repeated for each sample until the increase in number of reads aligning to the consensus sequences plateaued (2-9 iterations). After a final alignment of sRNA reads to the final consensus sequences for each sample, the total number of reads mapping to each virus was extracted from Geneious Prime. The ‘virus’ component of our sRNA composition (Figure 1) was calculated based on the total number of reads mapping to all viruses collectively after consensus strain iteration, divided by the total sRNA reads.

**Figure 1.**
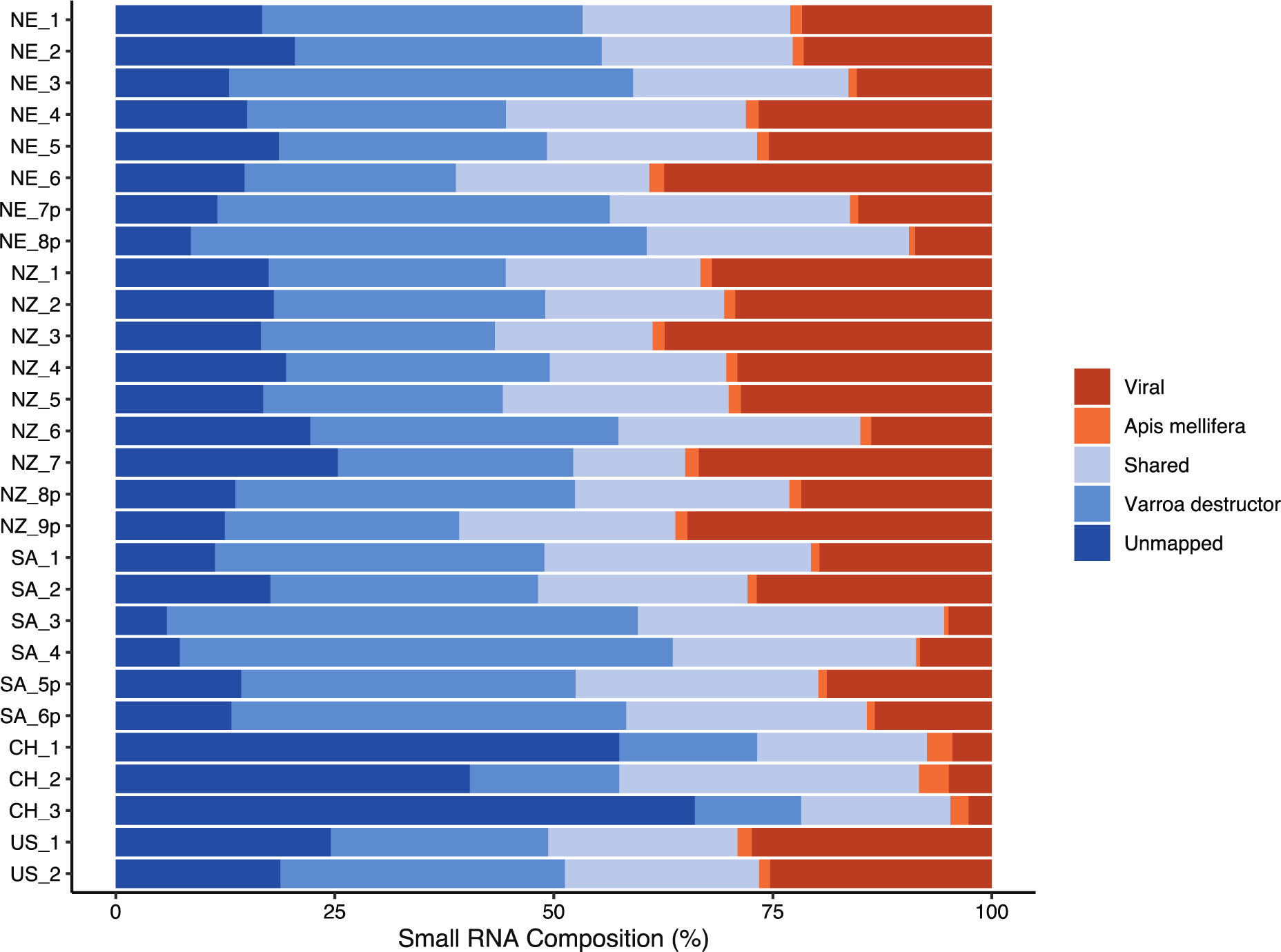
Small RNA composition of *V. destructor* samples. Each row indicates an individual mite sample; samples denoted ‘p’ are derived from four pooled mites. Country is indicated (NE = Netherlands; NZ = New Zealand; SA = South Africa; CH = China; US = United States). Reads were mapped to the *V. destructor* genome (Vdes_3.0; Genbank accession GCA_002443255.1) then the *A. mellifera* genome (Amel_HAv3.1; GCA_003254395.2); shared reads indicate those able to align to both species genomes. Viral reads represent the total vsiRNA reads aligned to all viruses identified in each sample after generation of sample-specific viral consensus genomes. The unmapped portion indicates reads that did not align to *V. destructor, A. mellifera* or examined viral consensus genomes.

### Phylogenetic analysis of novel and known viruses

The novel virus sequence segment identified in one of our Netherlands mite samples (NE-6) was used to probe previously published RNA sequencing data from 27 *V. destructor* samples from New Zealand (Lester et al., 2022) and two from Tonga (Remnant et al., 2017) to identify whether this novel virus was widespread and isolate a New Zealand variant. In addition, we used the RdRp-scan pipeline (Charon et al., 2022) and HMMER3 (Eddy, 2011) to search for the missing genomic segment of our putative novel virus containing the RdRp coding region.

To perform phylogenetic analysis of the Netherlands and New Zealand strains of our novel virus segment, we obtained homologous virus genomes from current NCBI databases using BLASTx, and performed protein alignments using MUSCLE (Edgar, 2004). We used IQ-TREE to generate maximum likelihood phylogenies with 1000 bootstraps and to determine the best-fit model (Nguyen et al., 2015). Treefiles were imported into FigTree (v1.4.4, http://tree.bio.ed.ac.uk/software/figtree/) and edited in Adobe Illustrator.

We also produced phylogenetic trees to examine nucleotide variation between global isolates of the viruses identified in our mite samples, using the strains assembled by Megahit or by consensus sequence generation after iterative alignments of the small RNA reads. After the final iteration of mapping, consensus sequences were generated for all viruses in each mite sample, leaving gaps over any regions which lacked coverage. Mite samples with lower virus levels returned consensus sequences that contained excessive gaps due to low coverage over some genomic regions and were thus excluded from further analysis. We constructed maximum likelihood phylogenetic trees using full length genomes for the most highly prevalent viruses (ARV-1, ARV-2, VDV-2, VDV-3/-5, DWV-A/-B), trimming regions with remaining short gaps in some samples. For DWV, we removed samples with evidence of recombinant variants, such as a DWV-B/A/B recombinant at the 5’ end of the genome in one of our Netherlands samples as previously characterised (Norton et al., 2020; Norton et al., 2021). Nucleotide sequences were aligned using MUSCLE, and IQ_TREE was used to generate a maximum likelihood phylogeny as described above.

### Analysis of vsiRNA

All sRNA reads were imported into Rstudio using Rsamtools. Read length distributions and genomic coverage from alignments to initial and curated viral reference library for single-mite samples were plotted in Rstudio using ggplot2 and custom scripts.

## RESULTS

### Small RNA composition in mites

Mite small RNA libraries comprised 12-56% of reads mapping to the *V. destructor* genome and 0.4-3.4% of reads mapping to *A. mellifera* (Figure 1). Due to the short nature of small RNA sequences and homology between genomes, 13-35% of reads mapped to both genomes, though we assume the majority of these reads are derived from *V. destructor*. The number of reads mapping to the viral reference library varied between mite samples. Visual inspection of vsiRNA alignments to the viral reference genomes could clearly show when a virus was present within a sample, however these alignments often had gaps, due to the highly polymorphic nature of RNA viruses, leading to siRNA reads with insufficient homology to viral reference genomes. To more accurately estimate the proportion of vsiRNA reads, consensus sequences for each virus were generated for each sample, and reads were re-aligned to the respective consensus genome sequences to capture the maximum number of viral reads. A significantly higher proportion of reads were found mapping to generated consensus genomes compared to the viral reference genomes (Figure S1). Across all samples, the final estimate of aligned viral reads ranged from 2.6-37% (Figure 1).

### Characterisation of a novel *V. destructor* virus

We performed *de novo* assembly of the small RNA reads to generate longer contigs to assist with producing sample-specific viral consensus strains (see Methods). Assembled contigs were examined for homology to known viruses using BLASTx against a database containing all available protein viral sequences (NCBI) to identify the sequences with homology to viruses. In three samples from the Netherlands (NE-6, 7p and -8p), we identified contigs that produced a full length 6.2 kb genome with 75% identity to Bee Macula-like virus (BMLV; NC_027631), indicating a novel variant (BMLV-NE 6-8; OR224322-OR224324; Figure S2 A). In addition, Netherlands sample (NE-6) we isolated two overlapping contigs that produced a 1.52 kb fragment with distant homology (31% amino acid identity) to the capsid protein of Beihai Horseshoe crab virus 1 (YP_009333375.1; Figure 2). Due to lack of homology to any *V. destructor* or *A. mellifera* sequences, or to any previously discovered viruses in bees or mites, we assumed that this sequence represented a segment of a novel virus. To determine if this novel virus was present in other mites, we additionally examined published *V. destructor* RNA-seq datasets from New Zealand (Lester et al., 2022) and Tonga (Remnant et al., 2017) and identified similar contigs with approximately 80% nucleotide identity to the novel virus in all transcriptomes examined (n = 27), suggesting this virus is common and widespread in *V. destructor*.

**Figure 2.**
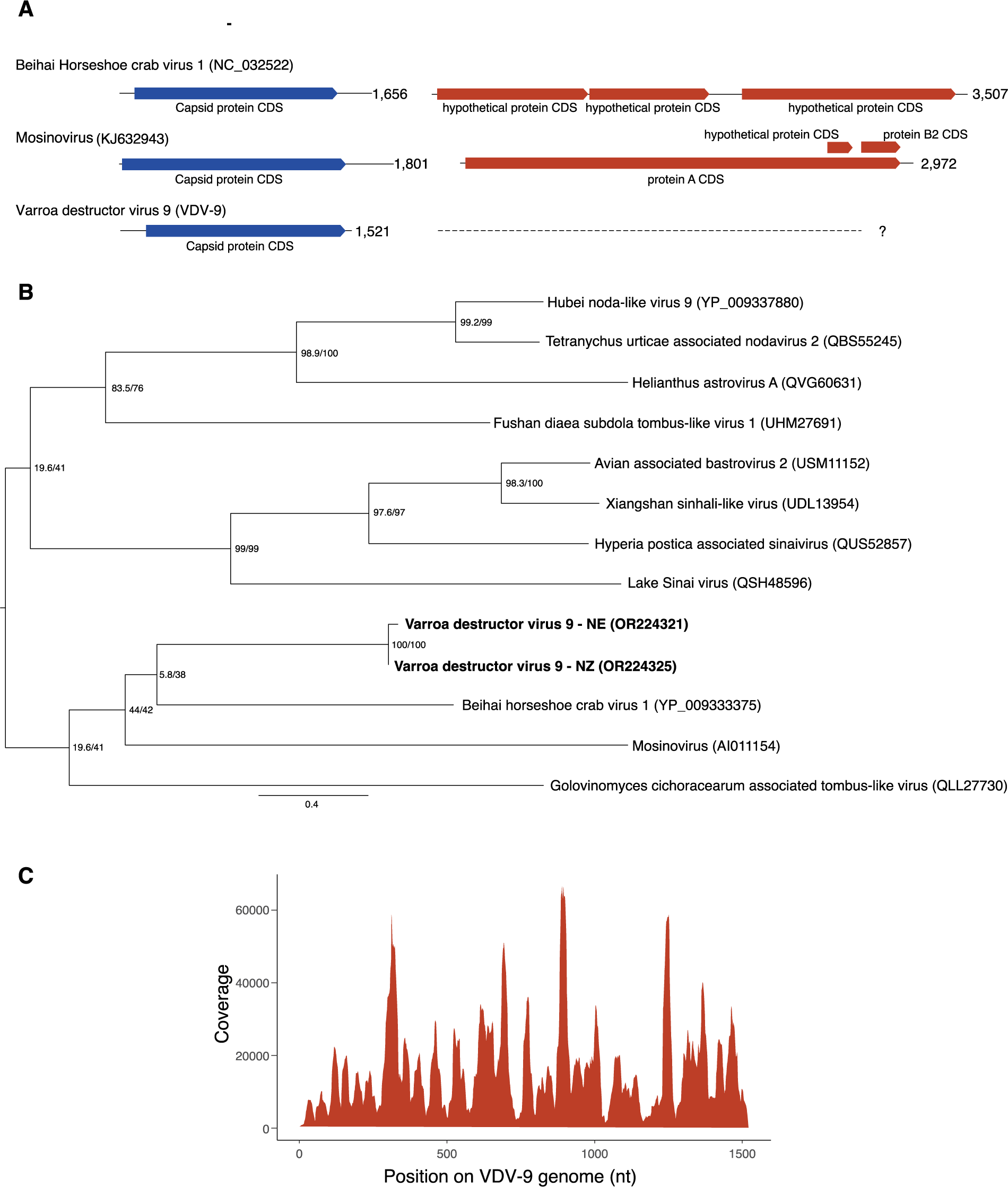
A) Putative genome structure of Varroa destructor virus 9 compared to two distantly related bi-segmented viruses, Behai horseshoe crab virus 1 and Mosinovirus. **B)** Phylogenetic analysis of the capsid protein segment of Varroa destructor virus 9 (VDV-9), identified in *V. destructor* samples from the Netherlands (NE; OR224321) and New Zealand (NZ; OR224325). Protein sequences of VDV-9 and homologous viruses obtained from NCBI were aligned using MUSCLE and trimmed for gaps, leaving a final length of 384 aa. The phylogenetic tree was generated using maximum likelihood in IQ-TREE (Nguyen et al., 2014), with the rtREV+F+I+G4 model which had the optimal BIC score, as determined by ModelFinder (Kalyaanamoorthy et al., 2017). Branch supports were estimated using Ultrafast bootstrap approximation (UFBoot; Hoang et al., 2017) using 1000 replicates. Support values shown are SH-aLRT and UFBoot supports. **C)** Genome coverage of small RNA reads mapped to VDV-9 (sample NE-6).

The distantly related Beihai Horseshoe crab virus 1 genome contains two positive-sense ssRNA subgenomic segments: a 1.6kb capsid segment with low homology to the fragment we identified in *V. destructor,* and a second 3.5kb segment containing the RNA dependent RNA polymerase (RdRp; Figure 2 A; Shi et al., 2016). We were unable to isolate the expected second segment of the *V. destructor* novel virus amongst our assembled contigs using BLASTx, or via HMMER3 using the RdRp-scan pipeline (Charon et al., 2022). As RNA viruses are usually classified based on their RdRp homology (Koonin et al., 2020), and we were only able to isolate the segment encoding the capsid protein in our novel virus, we were unable to assign a name that reflects taxonomy, so we named the novel virus ‘Varroa destructor virus 9’ (VDV-9; Genbank Accessions: OR224321 and OR224325). Phylogenetic analysis of the capsid protein sequence placed the Netherlands and New Zealand variants alongside Beihai Horseshoe crab virus 1 and Mosinovirus, isolated from mosquitoes (Schuster et al., 2014), with homology to Nodaviruses. Both Beihai Horseshoe crab virus 1 and Mosinovirus possess a bipartite, positive-sense RNA genome, with Mosinovirus containing an additional two sub-genomic segments (Schuster et al., 2014). The capsid protein of this clade is distantly related to members of the Tombus and Sinai virus family, including the honey-bee infecting Lake Sinai virus (Figure 2 B).

### vsiRNA composition in mites

We found evidence for a total of 10 viruses in mites across all of our samples, with the number of viruses present in individual mites ranging from three to nine viral species (Figure 3; Table 1). Positive-sense ssRNA viruses accounted for eight of the 10 viruses, along with two negative sense ssRNA viruses (ARV-1 and ARV-2). VDV-2 was the most prevalent virus, present in all *V. destructor* samples (n = 28), followed by ARV-1 and ARV-2 which were present in all but one sample from South Africa (SA-3). VDV-2 was also the most abundant virus, accounting for between 7-98% of vsiRNA reads, followed by DWV-A (0-89%) and DWV-B (0-39%). DWV abundance varied considerably between samples and was particularly high in four DWV-A containing mites from NZ (NZ-6, 7, 8p & 9p), accounting for 80-89% of viral reads.

**Figure 3:**
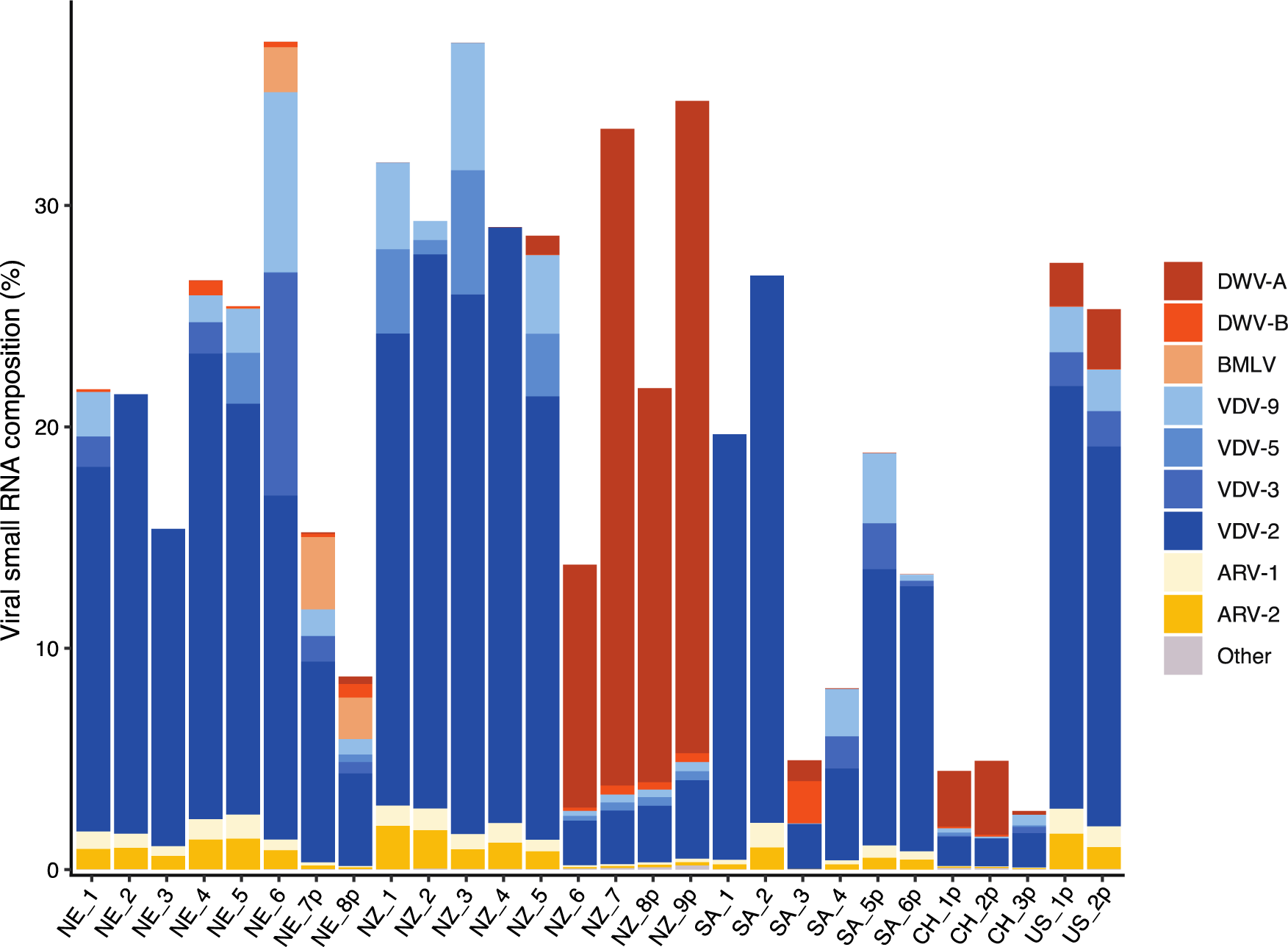
Composition of viruses within the viral small RNA portion of each sample. Each column represents an individual mite sample (named as in Figure 1). Y-axis indicates the percent of small RNA reads aligning to virus, with each colour indicating the proportion of total reads belonging to each virus species. ‘Other’ indicates the low proportion of reads aligning to viruses not listed; such as Sacbrood virus (SBV) and Black queen cell virus (BQCV) in NZ-9p. (ARV-1: Apis rhabdovirus 1; ARV-2: Apis rhabdovirus 2; VDV-2: Varroa destructor virus 2; VDV-3: Varroa destructor virus 3; VDV-5: Varroa destructor virus 5; VDV-9: Varroa destructor virus 9; BMLV: Bee macula-like virus; DWV-A: Deformed wing virus A; DWV-B: Deformed wing virus B).

**Table 1.**
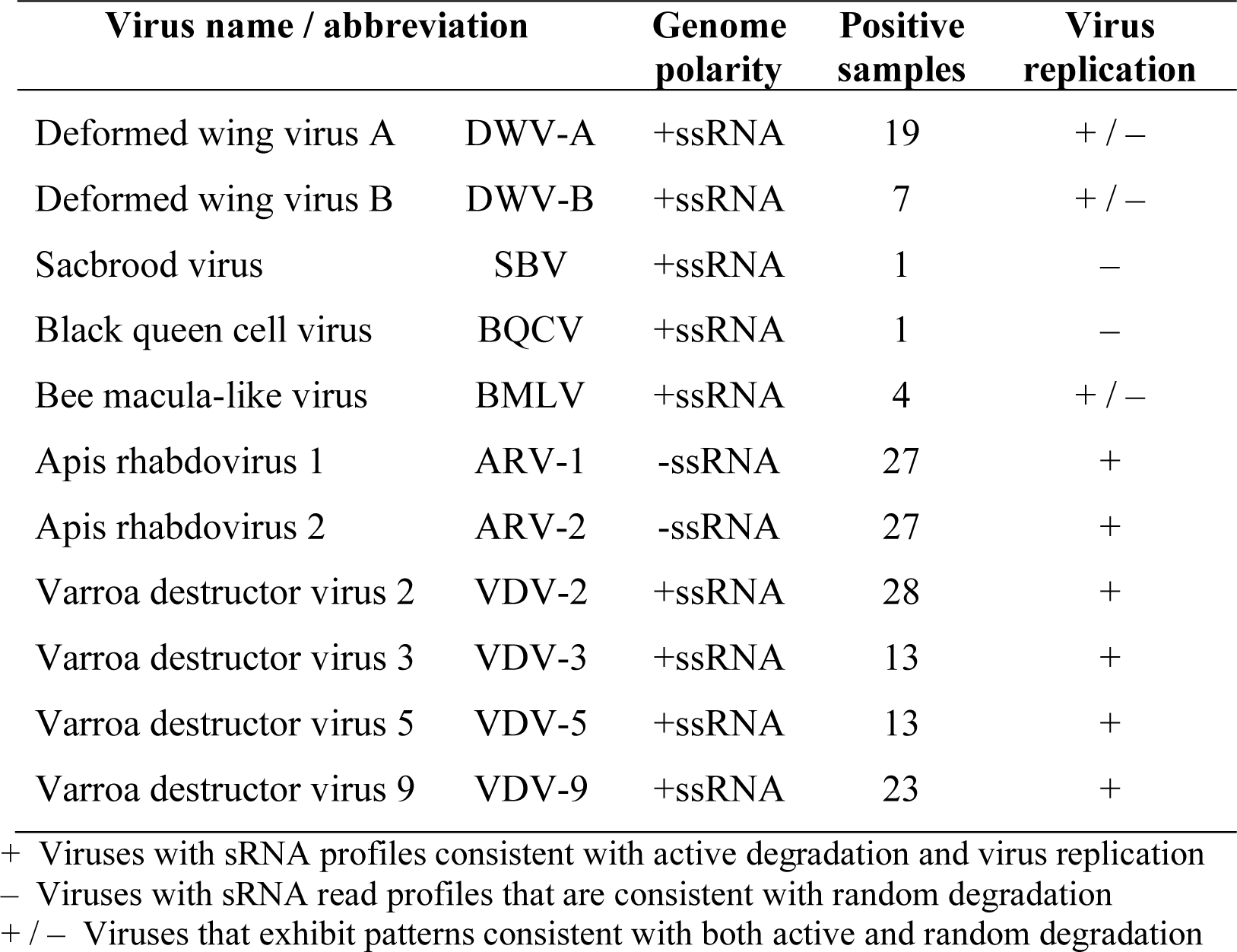
Summary of the replicating viruses identified in *Varroa destructor*.

| Virus name / abbreviation |  | Genome polarity | Positive samples | Virus replication |
| --- | --- | --- | --- | --- |
| Deformed wing virus A | DWV-A | +ssRNA | 19 | + / – |
| Deformed wing virus B | DWV-B | +ssRNA | 7 | + / – |
| Sacbrood virus | SBV | +ssRNA | 1 | – |
| Black queen cell virus | BQCV | +ssRNA | 1 | – |
| Bee macula-like virus | BMLV | +ssRNA | 4 | + / – |
| Apis rhadovirus 1 | ARV-1 | -ssRNA | 27 | + |
| Apis rhadovirus 2 | ARV-2 | -ssRNA | 27 | + |
| Varroa destructor virus 2 | VDV-2 | +ssRNA | 28 | + |
| Varroa destructor virus 3 | VDV-3 | +ssRNA | 13 | + |
| Varroa destructor virus 5 | VDV-5 | +ssRNA | 13 | + |
| Varroa destructor virus 9 | VDV-9 | +ssRNA | 23 | + |

+ Viruses with sRNA profiles consistent with active degradation and virus replication – Viruses with sRNA read profiles that are consistent with random degradation + / – Viruses that exhibit patterns consistent with both active and random degradation

### Phylogenetic clustering of consensus viral sequences

We performed phylogenetic analysis of the most prevalent viruses across individual samples (DWV-A/-B, Figure S2 B and ARV-1, ARV-2, VDV-2 and VDV-3/-5, Figure S3) by aligning the consensus sequences generated after iterative sRNA read mapping. Most viruses clustered broadly according to location with some individual exceptions (Figure S2 B and Figure S3). As VDV-3 and VDV-5 share approximately 75% nucleotide identity they were analysed in the same tree. The distributions of VDV-3 and -5 varied with NZ samples containing a VDV-5 variant (∼92% identity to VDV-5 reference), US and SA samples containing VDV-3, and the NE and CH samples showing evidence of both VDV-3 and VDV-5 variants (Figure S3 C).

Three separate VDV-2 consensus strains were generated for each sample, based on the three currently available reference genomes in NCBI (Israel, KX578271; UK, MK795517; and China, MW590582), which show between 75-83% nucleotide identity to each other. High coverage of all three reference strains in most samples (Supplementary file 1) indicates the presence of multiple VDV-2 strains within the same mite. Consensus sequence alignments clustered according to the reference VDV-2 sequence from which the consensus was originally based on, and then by location (Figure S3 D).

### vsiRNA profiles reveal active degradation of *Varroa-*infecting viruses

We examined the size profiles of viral small RNA reads mapping to all viruses present in our samples (Figure 4, Figure S4), including DWV-A and -B (Figure 5). The vsiRNA reads mapping to known *V. destructor*-infecting viruses (ARV-1 and -2, VDV-2, VDV-3 and VDV-5), have a clear antisense bias with 60-99% of reads in the antisense orientation, with some variation between viruses (Figure 4 A-E, Figure S4, Supplementary file 1). Most of the observed antisense reads have a size distribution peak at 24 nt, which is consistent between the -ssRNA viruses ARV-1 and -2 (Figure 4 A-B) and the +ssRNA viruses VDV-2, -3 and -5 (Figure 4 C-E). In addition to the 24 nt antisense bias, most viruses also have identifiable sense peaks at 20 and 23 nt, with the 20 nt peak more prominent in -ssRNA viruses and the 23nt peak more prominent in +ssRNA viruses. The novel virus, VDV-9 has an identical pattern to other *V. destructor*-infecting +ssRNA viruses with a strong 24 nt antisense peak, and a detectable 23 nt sense peak, indicating it is an actively replicating virus within *V. destructor* that can be degraded by the antiviral RNAi pathway (Figure 4 F).

**Figure 4:**
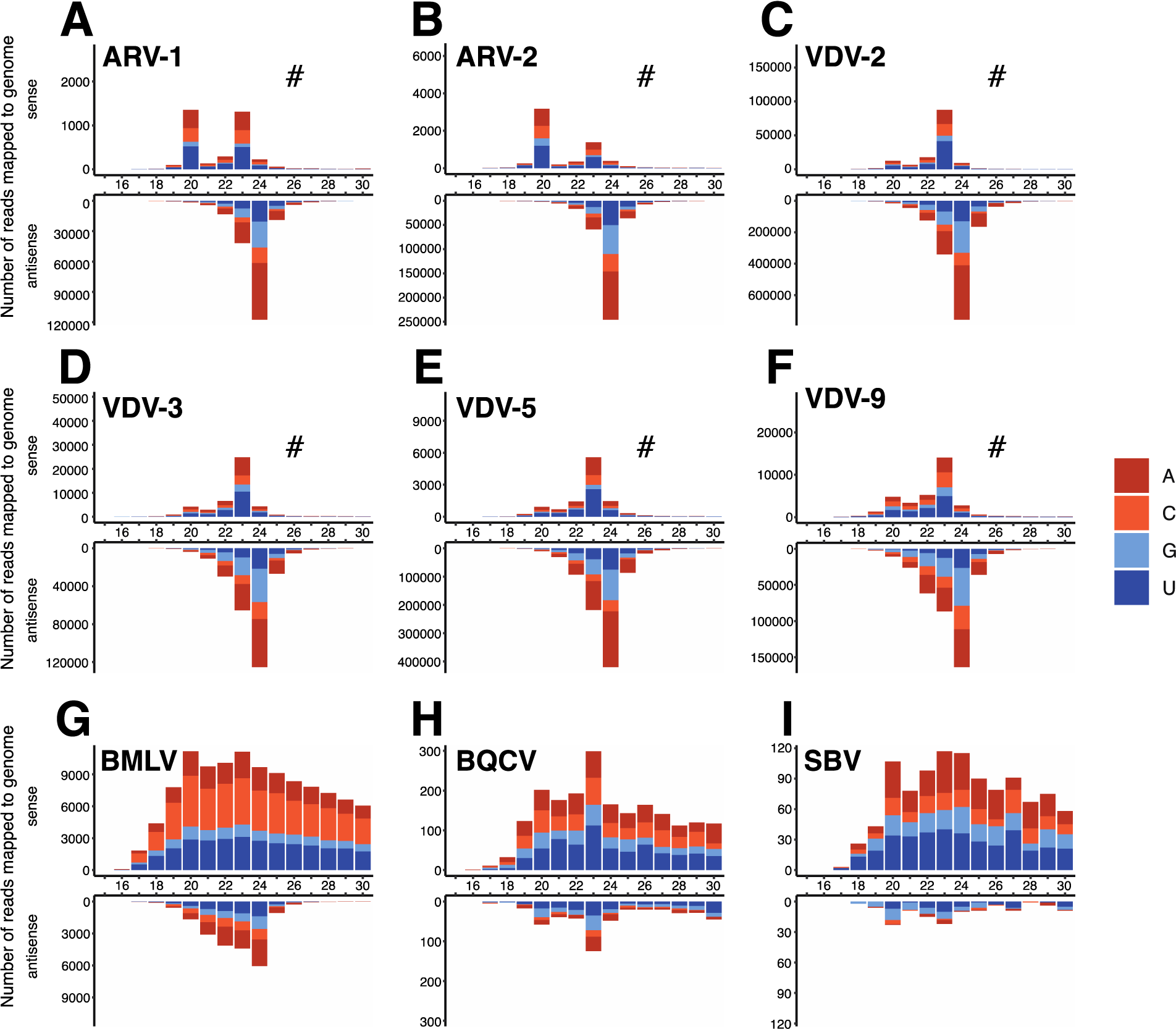
Size profiles of viral small RNA reads mapping to **A)** Apis rhabdovirus 1 (ARV-1); **B)** Apis rhabdovirus 2 (ARV-2); **C)** Varroa destructor virus 2 (VDV-2); **D)** Varroa destructor virus 3 (VDV-3); **E)** Varroa destructor virus 5 (VDV-5); **F)** Varroa destructor virus 9 (VDV-9); **G)** Bee macula-like virus (BMLV); **H)** Black queen cell virus (BQCV); and **I)** Sacbrood virus (SBV). Representative profiles are shown from samples chosen at random (NZ-4: A, C; NZ-1, B; US-1p: D, F; NZ-3: E; NE-8p: G; NZ-9p: H, I) with near identical patterns observed for all mites infected with the above viruses (see Figure S4 for additional examples). # indicates sense Y-axis has been scaled to aid in visualising any size peaks.

**Figure 5.**
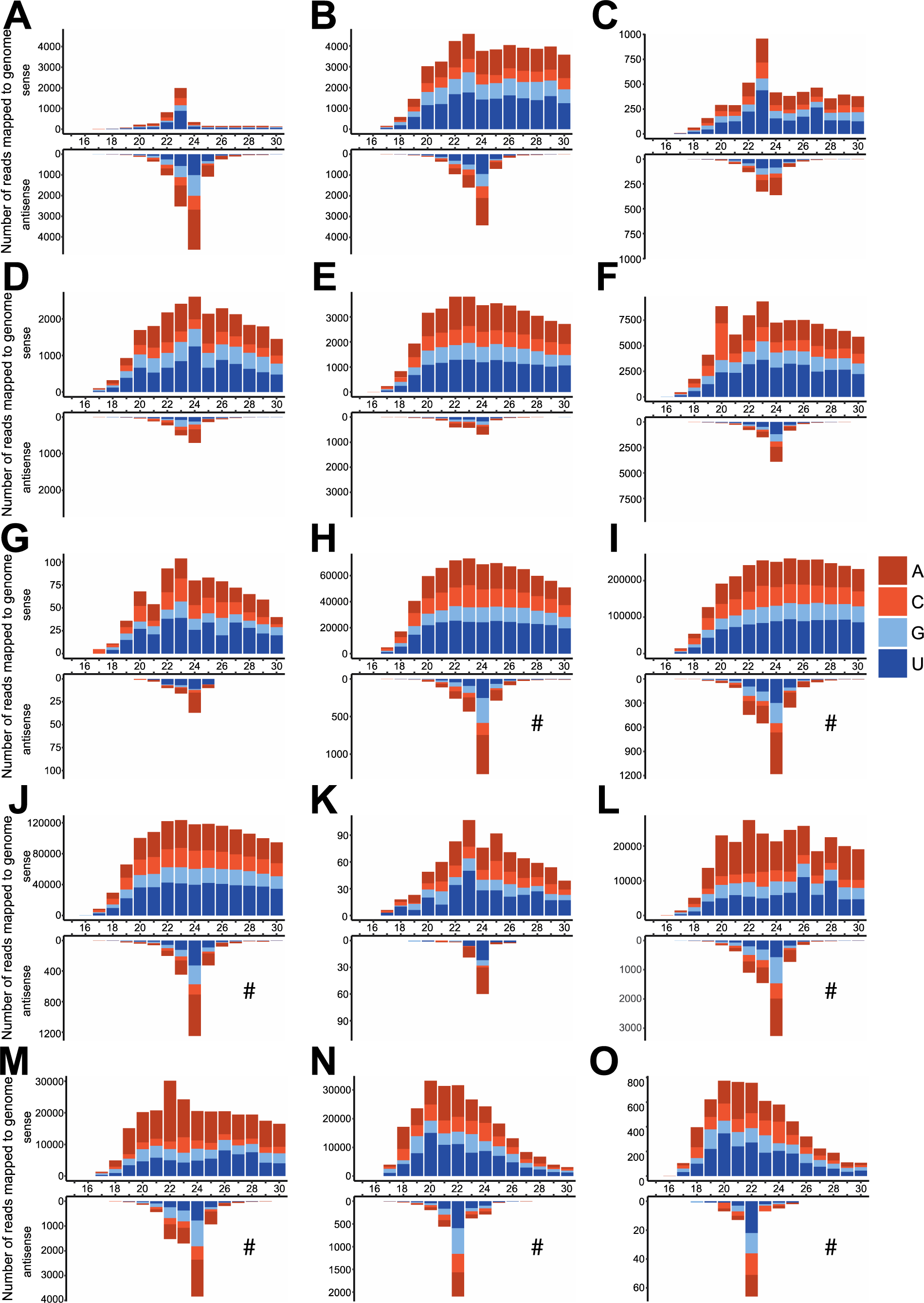
Size profiles of viral small RNA reads mapping to DWV-B (A-D) and DWV-A (E-O) from **A)** NE-1; **B)** NE-4; **C)** NE-5; **D)** NE-6; **E)** NE-8p; **F)** NZ-5; **G)** NZ-2; **H)** NZ-6; **I)** NZ-7; **J)** NZ-8p; **K)** SA-5p; **L)** US-1p; **M)** US-2p, **N)** CH-2p; **O)** CH-3p. # indicates antisense Y-axis has been scaled to aid in visualising any size peaks.

Honey bee-associated viruses (BMLV, BQCV and SBV) have more ambiguous degradation profiles. BMLV has a mixed profile with an antisense 24 nt peak, though the majority of mapped reads appear to be passively degraded according to the high proportion of sense reads (27-84%) which correspond to the genome polarity of BMLV (+ssRNA; Figure 4 G, Figure S4 G, Supplementary file 1). BQCV and SBV were identified in one pooled mite sample (NZ-9p), but these viruses are in low abundance, lack a 24 nt antisense peak, and have a high proportion of sense reads (>80%), consistent with passive degradation (Figure 4 H-I).

While most *V. destructor*-infecting viruses display consistent active degradation profiles between all mite samples (Figure 4 A-F and Figure S4), DWV shows variable profiles between individual mites (Figure 5). In one DWV-B-infected sample from the Netherlands (NE-1) there is a large proportion of antisense vsiRNA reads (69%) with a clear 24 nt peak and an additional 23nt sense peak (Figure 5 A), similar to the active degradation profiles of other +ssRNA viruses (Figure 4 C-F). Other DWV-infected mites have evidence of a 24 nt antisense peak along with a 23 nt sense peak (eg. Figure 5 C-F), though with a higher proportion of sense reads. In particular, in samples where DWV vsiRNA reads are highly abundant (Figure 3), the majority of reads aligning to DWV have a positive sense distribution consistent with DWV genome polarity (+ssRNA), and a much lower proportion of antisense reads, though with an observable antisense peak at 24 nt in most samples (Figure 5 H-J, L-N). Interestingly, two mite samples from China (CH-2p and CH-3p, Figure 5 N, O) show a 22 nt antisense peak, consistent with the mites having ingested honey bee-produced vsiRNAs (Chejanovsky et al., 2014; Remnant et al., 2017). Collectively, the degradation profiles of DWV-A and -B genotypes indicate a mixed mode of active and passive degradation, suggesting that both genotypes have evidence of active replication in mites, but that a considerable viral load is also passively degraded, either due to ingested honey bee tissues, or due to unchecked viral replication and escape from RNAi control within mite cells (Figure 5).

### Replicating viruses in *V. destructor*

Based on the active vsiRNA profiles identified, we summarise the replication status of all viruses present in our *V. destructor* samples (Table 1). A total of eight viruses (including both DWV-A/- B strains) have active degradation profiles indicative of viral replication in *V. destructor*.

## DISCUSSION

In this study, we identified eight actively replicating viruses in *V. destructor,* based on host-specific vsiRNA profiles obtained by examining small RNA from mites sampled globally. Most notably, we provide direct evidence for replication of DWV-A and -B within mites, suggesting that *Varroa* is a biological vector of both genotypes. Our analysis of vsiRNA profiles in *V. destructor* identifies clear antiviral degradation with both +ssRNA and -ssRNA viruses showing a 24 nt antisense peak, and additional minor sense peaks. A mixed active/passive degradation pattern observed in some DWV samples was also evident in another honey-bee associated virus, BMLV, but not SBV and BQCV. We also present evidence of a novel, bi-segmented +ssRNA virus that is widespread in *V. destructor*, VDV-9, that shows a 24 nt vsiRNA degradation profile and is therefore likely to be an actively replicating virus in mites.

*V. destructor* is associated with an array of viruses, many of which are due to its role as a parasite of *A. mellifera* and viral vector of bee-infecting viruses. Due to the critical role of *V. destructor* in the transmission of bee viruses, it is important to understand the mechanisms in which mites achieve antiviral immunity, and to identify viruses that are replicating within mites. Identification of replicating +ssRNA viruses in honey bees and other invertebrates is normally achieved via strand-specific RT-PCR of negative sense strands, which are produced during viral replication (Ongus et al., 2004; Runckel et al., 2011; Yue and Genersch, 2005). A limitation of using such methods for parasites like *Varroa* is that they ingest infected host tissues that could contain negative sense replication intermediates, which will give false positive signals. However, since *A. mellifera* and *V. destructor* have different antiviral siRNA profiles (Remnant et al., 2017), we took advantage of these differences to comprehensively characterise vsiRNA profiles in individual mites to identify replicating viruses and determine whether *V. destructor* is a genuine host of these viruses. For most viruses present in our mite samples, we observed vsiRNA profiles with a 24 nt antisense bias (Figure 4). Polarity biases during RNAi-mediated antiviral defence are not unique to *V. destructor*, as similar polarity biases have been observed in *C. elegans* (Ashe et al., 2013), *Aedes aegypti* (Brackney et al., 2009), and a range of other metazoans (Sekhar Nandety et al., 2013; Waldron et al., 2018). We categorised viruses with a distinct 24 nt antisense peak as replicating viruses within *V. destructor,* and identified eight viruses replicating within our samples based on this criteria (Table 1). Notably, all *V. destructor* samples contained VDV-2 and all but one sample had ARV-1 and ARV-2 (Figure 3; Table 1), which are three viruses known to be highly prevalent in *V. destructor* (Eliash et al., 2022; Herrero et al., 2019; Lester et al., 2022; Levin et al., 2016). We saw evidence of individual mites containing multiple VDV-2 strains (Figure S3 D), as previously observed in New Zealand *V. destructor* samples (Lester et al., 2022), though this is difficult to resolve conclusively using small RNA data, due to short read lengths and nucleotide similarities between strains. Based on the high prevalence of ARV-1 and ARV-2 in mites, and low prevalence in honey bees and other bee species (Levin et al., 2017; Norton et al., 2021) there is accumulating evidence that *V. destructor* is the primary host for these -ssRNA rhabdoviruses instead of *A. mellifera*, and mounting evidence that these viruses frequently co-occur in both bees (Kadlečková et al., 2022) and mites. Additional Rhabodviruses have also been identified in *Varroa jacobsoni* (Roberts et al., 2020), as well as within *A. mellifera* and *Apis cerana* in China (Li et al., 2023), indicating substantial diversity in the -ssRNA rhabovirus clade.

The novel viral genome identified in this study, Varroa destructor virus 9 (VDV-9), was also present in 23 (82%) of our mite samples (Figure 3, Table 1), with vsiRNA profiles consistent with viral replication (Figure 4 F). The genomic segment identified in our samples has low homology to the capsid protein of other +ssRNA bi-segmented viruses including Behai Horseshoe crab virus 1 and Mosinoviruses (Figure 2), yet we were unable to isolate the RdRp-coding segment in our data. It is therefore likely that a portion of the ‘unmapped’ small RNA reads (Figure 1) belong to our as yet unidentified RdRp-containing fragment, or indeed to other as yet unidentified viruses in these samples. It is possible that the predicted VDV-9 segment containing the RdRp coding region lacks sufficient homology to currently known viruses and therefore cannot be identified by available methods. Identifying highly divergent viral sequences from the ‘dusk matter’ of metatranscriptomic datasets is a common challenge, and virus-derived small RNA has been successful in identifying novel lineages that lack sufficient homology to known viruses (Obbard et al., 2020; Webster et al., 2015). However, our RdRp-specific HMM search, which can identify homology as low as 10% (Charon et al., 2022) did not yield any novel contigs with RdRp homology from our small RNA samples or additional transcriptomes from New Zealand *V. destructor*. Alternatively, the capsid segment we identified here might represent the outcomes of a novel viral reassortment (McDonald et al., 2016), where this segment has become associated with another, already-known virus present in mites. The vast majority of viruses identified in *Varroa* have been characterised at sequence level only, with no knowledge of viral particle structure or understanding of disease phenotypes, thus it is not unfeasable that viral segments may be missing from previously characterised viruses. There is clearly more to be done to understand the role of VDV-9 as a member of the *V. destructor* virome. More broadly, we require a deeper understanding of how the diverse and complex virome present in individual mites, which contain between 3-9 viruses, impacts *V. destructor* biology.

Although BQCV and SBV are commonly among the most prevalent viruses identified in honey bee samples globally (Bailey, 1967; Li et al., 2023; Mondet et al., 2014; Roberts et al., 2017; Tentcheva et al., 2004), they were detected in only one of our *V. destructor* samples (NZ-9p). Negative strand RT-PCR assays for BQCV has previously implied active replication occurs in *V. destructor,* suggesting it could be a biological vector for this virus (Sabahi et al., 2020); however is possible that BQCV negative strand RT-PCR may have detected honey bee infected tissue within *V. destructor.* From our limited availability of positive samples (n=1), the distribution of small RNA reads mapping to BQCV is not consistent with replication and shows a random degradation profile (Figure 4 H). Similarly, SBV vsiRNA read profiles were also inconsistent with replication (Figure 4 I), suggesting that *V. destructor* is not a biological vector of BQCV or SBV, though it may still transmit viral particles passively as a mechanical vector. Analysis of small RNA profiles of additional mites containing BQCV and SBV is necessary before this can be conclusively stated. The lack of *V. destructor* samples with BQCV and SBV supports the hypothesis that viruses such as these, which show high lethality in juvenile life stages (Bailey, 1969; Bailey and Woods, 1977) are incompatible with the *V. destructor* life cycle, leading to selection against vectoring by mites (Martin, 2001; Remnant et al., 2019).

Because DWV is the main virus associated with colony declines, the question of whether *V. destructor* is a biological vector and a true host for DWV has been a topic of great interest in honey bee research (Beaurepaire et al., 2020; Gisder et al., 2009; Piou et al., 2022; Sabahi et al., 2020). To add to growing experimental data that indicate active replication of DWV-B and recombinant DWV genotypes within *Varroa* tissues (Gisder and Genersch, 2021; Gusachenko et al., 2020), we provide direct evidence of DWV-A and -B replication in *V. destructor* via the presence of active antiviral degradation profiles.

Multiple DWV-infected mites show vsiRNA profiles with distinct 24 nt antisense peaks (Figure 5), as seen in actively replicating +ssRNA viruses in *V. destructor* (Figure 4 C-F), indicating that there is active degradation and therefore viral replication of DWV within mites. This active degradation profile was present in mite samples containing DWV-A and -B (Figure 5 A-M). Some samples show a mixed profile with the 24 nt antisense peak and a higher proportion of sense reads showing random degradation, which could indicate that individual mites have low levels of DWV replication, and the vsiRNA signal is overwhelmed by the high proportion of ingested DWV from infected *A. mellifera* tissue, resulting in a random degradation profile that swamps the proportion of antisense reads. Mites with lower DWV loads tended to have clearer active degradation profiles, in support of this hypothesis. It is likely that the use of whole crushed mites yielded higher positive-sense passively degraded reads, because small RNA from all tissues was represented, including from the digestive tract which contains degradation products of ingested viruses. Alternatively, the high positive sense reads in some samples could indicate that actively replicating DWV virus has overwhelmed the RNAi response exceeded their immune capacity to achieve sufficient viral control. Of note, two samples (Figure 5 N-O) show clear 22 nt antisense peaks, evidence that these mites have ingested honey bee-produced vsiRNAs rather than generated their own antiviral response. This outcome shows the sensitivity of our method in detecting *V. destructor*-generate antiviral small RNA fragments, which has a clear size difference (24 nt) to *A. mellifera-*generated fragments (22 nt).

*In situ* hybridisation shows that replicating DWV-B is localised to gut epithelial tissue and salivary glands, reinforcing the notion that DWV replicates in key tissues of *V. destructor* that are compatible with biological vectoring (Gisder and Genersch, 2021). In the same study, DWV-A replication was not detected by *in situ* hybridisation, though the authors did not confirm the presence of DWV-A in their source population. Our data suggests that there is no strain specificity regarding DWV replication. While individual mite DWV vsiRNA profiles vary considerably along with variation in DWV loads, overall, the presence of some degree of active antiviral response to DWV-A (Figure 5 E-M) and DWV-B (Figure 5 A-D) in multiple individual mites indicates that *V. destructor* is a genuine host for both DWV-A and -B genotypes.

Currently, the mechanism underpinning the 24 nt antisense peak that differentiates the *V. destructor* vsiRNA profile from the typical 21-22 nt sense/antisense profile observed honey bees and other insects is unknown, however it draws parallels to the secondary amplification of siRNA by an RNA-directed-RNA-polymerase (RdRp) as found in the roundworm, *Caenorhabditis elegans* (Ashe et al., 2013). Secondary amplification of exogenous siRNA by RdRps in *C. elegans* results in synthesis of abundant antisense siRNAs that are distinct from primary siRNAs produced by the Dicer pathway. In plants and *C. elegans*, the production of secondary siRNA enhances antiviral immunity. Notably however, the size profiles of secondary siRNAs in *C. elegans* are 22 nt (Akay et al., 2015; Ashe et al., 2013). RdRp orthologs are largely absent from most insects including honey bees (Pinzón et al., 2019), however they have been identified in members of the Acari subclass (ticks and mites), with four identified in the *V. destructor* genome (Nganso et al., 2020). The role of RdRps during RNAi-mediated antiviral defence has not been studied in *Varroa*, but some work has been done within other members of Acari. Like *V. destructor,* the Black-legged tick *Ixodes scapularis* has four RdRps within its genome. However in *I. scapularis*, RNAi-mediated antiviral immunity generates vsiRNAs which are 22 nt in length, with no clear polarity bias (Schnettler et al., 2014) suggesting that RdRps are unlikely to be involved in secondary siRNA synthesis in this species. The *V. destructor* genome contains multiple Dicer-2 and Ago-2 isoforms (Nganso et al., 2020), and gene expansions in these other components involved siRNA-mediated antiviral immunity may have functionally distinct roles in mites. It is possible that the presence of multiple *Dicer-2* homologues in the *V. destructor* genome result in different small RNA populations in a similar mechanism to plant antiviral defence, where 21, 24, and 22 nt siRNA have been observed to be generated by different *Dicer* genes (Borges and Martienssen, 2015; Waterhouse and Fusaro, 2006). Currently, it is unclear which of the RNAi pathway components produces *V. destructor*’s uniquely sized antisense siRNA profile in response to viral replication, or if mite RdRps are used to enhance antiviral immunity in a similar mechanism to *C. elegans*. Nevertheless, the alternative antiviral RNAi mechanism present in mites has allowed us to successfully identify replicating viruses within *V. destructor*, most notably with DWV.

Our study shows that *V. destructor* harbours a core virome that is abundant and diverse, which is actively targeted by the mite’s antiviral RNAi pathway. Our study has added to this diversity with identification of a novel, capsid-encoding segment of a putatively bi-segmented +ssRNA virus, VDV-9, with homology to Tombus and Sinai-like viruses (Figure 2). It is presently unclear whether mite-infecting viruses cause pathology in *V. destructor*. Enhanced antiviral protection may be required to protect mites against their own viruses, or to withstand the pathogenic impact of the viruses that mites vector to honey bees. Our results suggest that the role of *V. destructor* in virus transmission is mixed, and involves both active and passive transmission. Ultimately our results indicate that honey bee viruses like DWV-A, DWV-B and BMLV have adapted to replicate within mites, as a result of viral spillover from the honey bee host. Host-parasite viral spillover is therefore a key contributing factor to the increasing impact of viruses on honey bee health as a result of the global spread of *V. destructor*.

## Acknowledgements

We thank Phil Lester, Madeleine Beekman and Ben Oldroyd for assistance with sample collection, and Julie Lim for laboratory and moral support. We acknowledge the Sydney Informatics Hub and The University of Sydney’s high performance computing cluster Artemis for providing the high-performance computing resources that have contributed to the research results reported in this paper.

