## Supplementary Tables and Figures for "Virus replication in the honey bee parasite, *Varroa destructor*"

**Table S1.** *Varroa destructor* small RNA samples used in this study.

**Figure S1.** Comparison of the proportion of small RNA reads mapping to virus genomes before and after consensus sequence generation

**Figure S2.** Phylogenetic analysis of honey bee-associated viruses in *V. destructor* samples.

**Figure S3.** Phylogenetic analysis of consensus sequences of *V. destructor*-associated viruses.

**Figure S4.** Additional examples of size profiles of viral small RNA reads mapping to viruses

**Supplementary file 1.** Spreadsheet showing the number and proportion of sense and antisense reads mapping to the viruses identified in this study:

- ARV-1
- ARV-2
- VDV-2
- VDV-3
- VDV-5
- VDV-9
- DWV-A
- DWV-B
- BMLV
- BQCV
- SBV

**Table S1.** *Varroa destructor* small RNA samples used in this study. All samples are individual whole mites other than those denoted with p, which consist of pooled mites. In our samples we pooled 4 mites, while other studies pooled 20 (CH-1p, CH-2p, CH-3p, US-1p) or 10 mites (US-2p).

| Sample | Location | Collection date | Biosample ID (SRA) | Reference |
|---|---|---|---|---|
| NE-1 | Wageningen, Netherlands | Aug 2018 | SRR25010770 | *This study* |
| NE-2 | Wageningen, Netherlands | Aug 2018 | SRR25010769 | *This study* |
| NE-3 | Wageningen, Netherlands | Aug 2018 | SRR25010758 | *This study* |
| NE-4 | Wageningen, Netherlands | Sep 2018 | SRR25010756 | *This study* |
| NE-5 | Wageningen, Netherlands | Sep 2018 | SRR25010755 | *This study* |
| NE-6 | Wageningen, Netherlands | Sep 2018 | SRR25010754 | *This study* |
| NE-7p | Wageningen, Netherlands | Aug 2018 | SRR25010753 | *This study* |
| NE-8p | Wageningen, Netherlands | Aug 2018 | SRR25010752 | *This study* |
| NZ-1 | Wellington, New Zealand | Mar 2022 | SRR25010751 | *This study* |
| NZ-2 | Wellington, New Zealand | Mar 2022 | SRR25010750 | *This study* |
| NZ-3 | Wellington, New Zealand | Mar 2022 | SRR25010768 | *This study* |
| NZ-4 | Wellington, New Zealand | Mar 2022 | SRR25010767 | *This study* |
| NZ-5 | Hamilton, New Zealand | Mar 2015 | SRR25010766 | *This study* |
| NZ-6 | Wellington, New Zealand | Mar 2022 | SRR25010765 | *This study* |
| NZ-7 | Wellington, New Zealand | Mar 2022 | SRR25010764 | *This study* |
| NZ-8p | Wellington, New Zealand | Mar 2022 | SRR25010763 | *This study* |
| NZ-9p | Wellington, New Zealand | Mar 2022 | SRR25010762 | *This study* |
| SA-1 | Robben Island, South Africa | Mar 2013 | SRR5109825 | (1) |
| SA-2 | Robben Island, South Africa | Mar 2013 | SRR5109827 | (1) |
| SA-3 | Kromme Rhee, South Africa | Nov 2016 | SRR25010761 | *This study* |
| SA-4 | Kromme Rhee, South Africa | Nov 2016 | SRR25010760 | *This study* |
| SA-5p | Elsenburg, South Africa | Nov 2016 | SRR25010759 | *This study* |
| SA-6p | Elsenburg, South Africa | Nov 2016 | SRR25010757 | *This study* |
| CH-1p | Jilin, China | Oct 2018 | SRR13959567 | (2) |
| CH-2p | Heilongjian, China | Oct 2018 | SRR13959568 | (2) |
| CH-3p | Hebei, China | Nov 2018 | SRR13959569 | (2) |
| US-1p | Mississippi, USA | Oct 2019 | SRR17429192 | (3) |
| US-2p | Mississippi, USA | Oct 2019 | SRR17429196 | (3) |

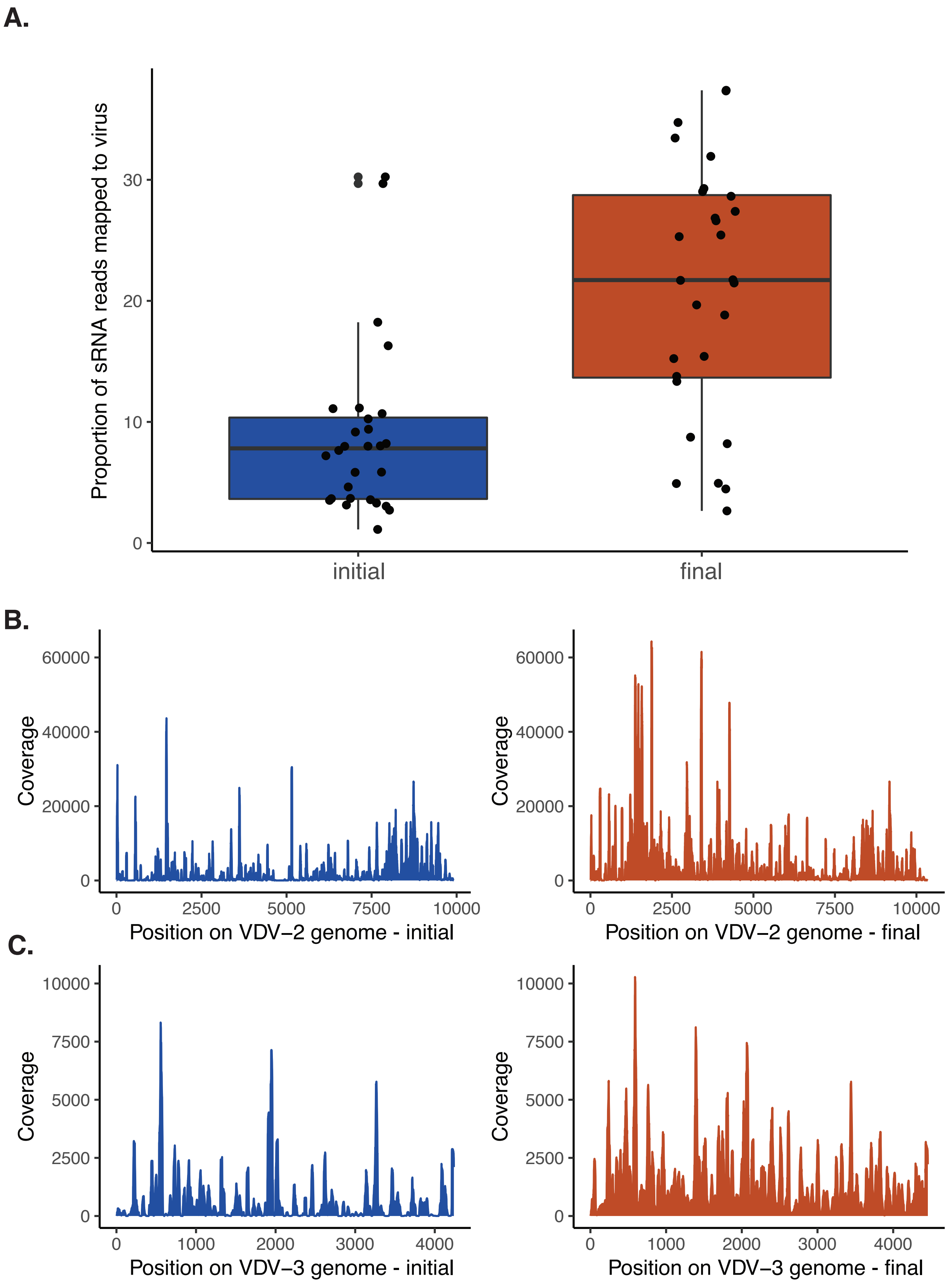


**Figure S1. A)** Comparison of the proportion of small RNA reads mapping to reference virus genomes (initial; blue) versus the proportion of reads mapping to consensus genomes generated for each virus after iterative mapping (final; orange). **B-C)** Coverage plots of **B)**VDV-2 and **C)** VDV-3 genomes of reads mapping to virus reference genomes (initial; blue) versus coverage of reads mapping to consensus genomes generated after iterative mapping (final; orange), showing improved coverage and reduced regions with gaps. Representative samples are shown (**B)** VDV-2: NZ-3; **C)** VDV-3: SA_5p).

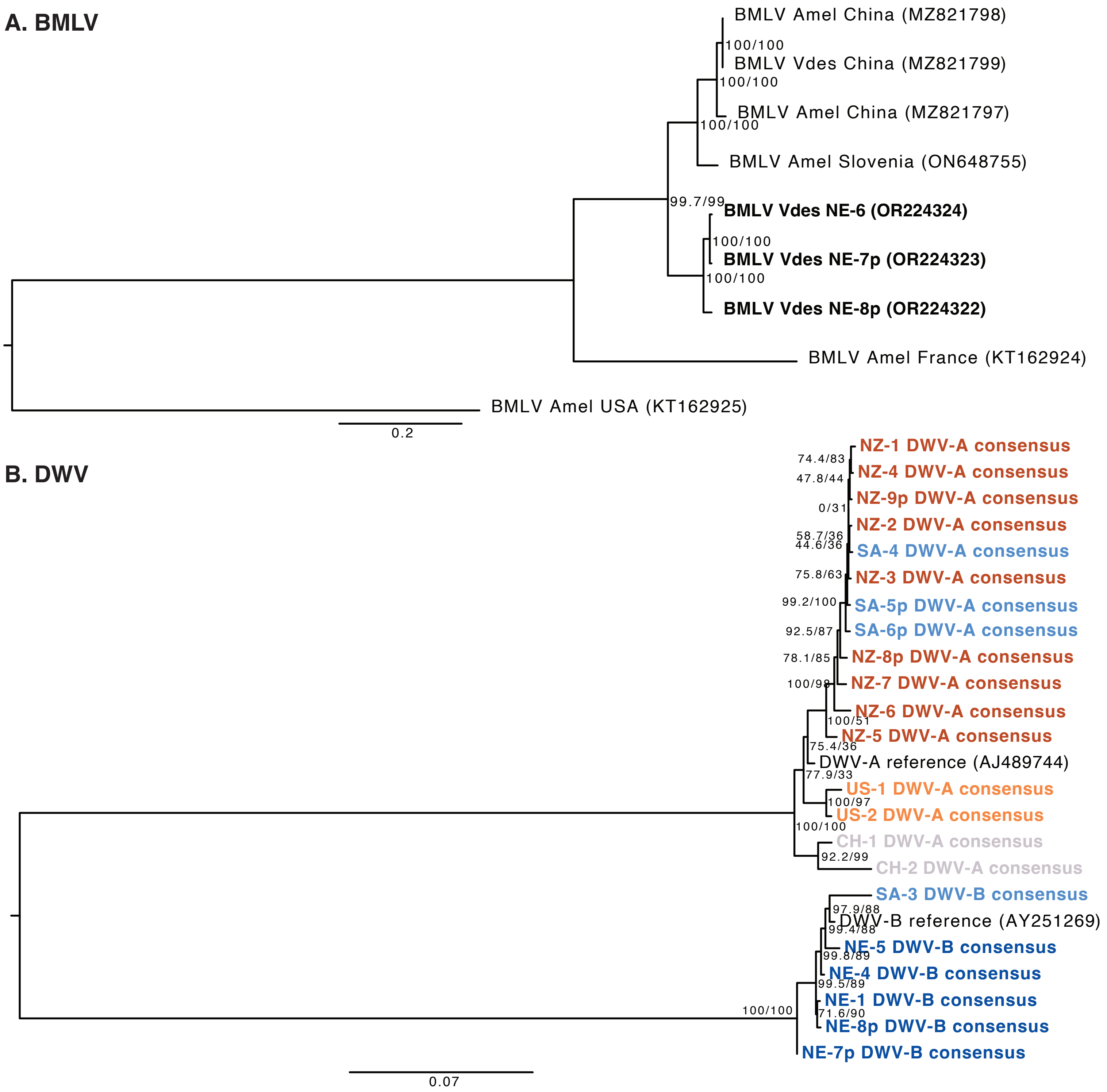


**Figure S2.** Phylogenetic analysis of honey bee- associated viruses in *V. destructor* samples. **A)** Bee macula-like virus isolates from three *V. destructor* samples from the Netherlands (OR224322-OR224324). Whole genome nucleotide sequence alignments were aligned using MUSCLE. After trimming, alignment length was 5930 nt. The phylogenetic tree was generated using maximum likelihood in IQ-TREE [1], with the TN+F+I+G4 model which had the optimal BIC score, as determined by ModelFinder [2]. **B)** Deformed wing virus consensus sequences from positive *V. destructor* samples. Whole genome nucleotide sequences were aligned as above and samples with evidence of recombination were removed. The final length after trimming was 9198 nt. Phylogenetic tree generation and model prediction were performed as above, with the TN+F+G4 model chosen according to BIC. For both A and B, branch supports were estimated using Ultrafast bootstrap approximation (UFBoot; [3]) using 1000 replicates. Support values shown are SH-aLRT and UFBoot supports.

**1.** Nguyen, L.-T.; Schmidt, H.A.; von Haeseler, A.; Minh, B.Q. Iq-tree: A fast and effective stochastic algorithm for estimating maximum-likelihood phylogenies. Mol Biol Evol 2014, 32, 268-274.
**2.** Kalyaanamoorthy, S.; Minh, B.Q.; Wong, T.K.F.; von Haeseler, A.; Jermiin, L.S. Modelfinder: Fast model selection for accurate phylogenetic estimates. Nat Methods 2017, 14.
**3.** Hoang, D.T.; Chernomor, O.; von Haeseler, A.; Minh, B.Q.; Vinh, L.S. Ufboot2: Improving the ultrafast bootstrap approximation. Mol Biol Evol 2017, 35, 518-522.

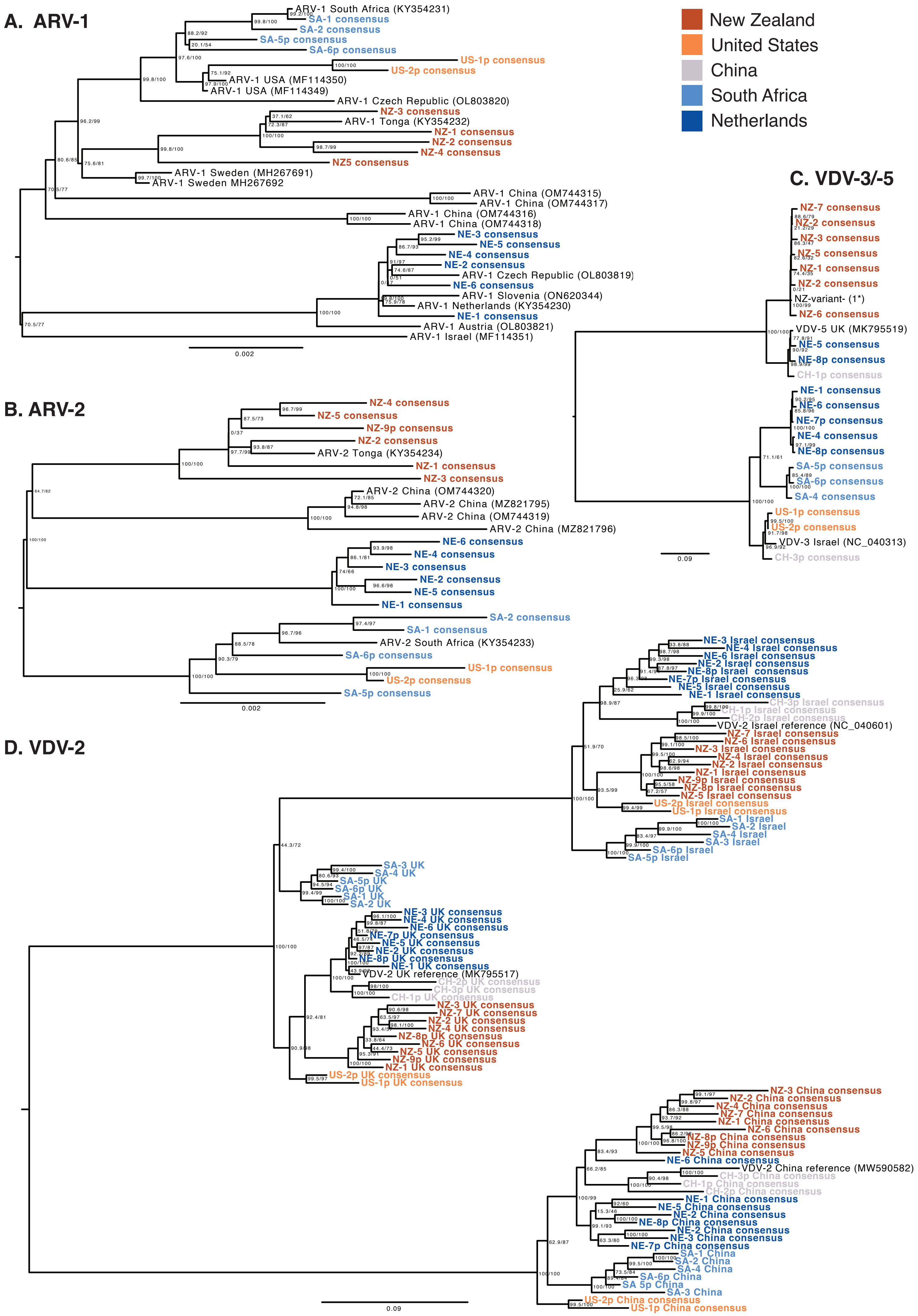

A. ARV-1
ARV-1 South Africa (KY354231)
SA-1 consensus
SA-2 consensus
SA-5p consensus
SA-6p consensus
US-1p consensus
US-2p consensus
ARV-1 USA (MF114350)
ARV-1 USA (MF114349)
ARV-1 Czech Republic (OL803820)
NZ-3 consensus
ARV-1 Tonga (KY354232)
NZ-1 consensus
NZ-2 consensus
NZ-4 consensus
NZ5 consensus
ARV-1 Sweden (MH267691)
ARV-1 Sweden MH267692
ARV-1 China (OM744315)
ARV-1 China (OM744317)
ARV-1 China (OM744316)
ARV-1 China (OM744318)
NE-3 consensus
NE-5 consensus
NE-4 consensus
NE-2 consensus
ARV-1 Czech Republic (OL803819)
NE-6 consensus
ARV-1 Slovenia (ON620344)
ARV-1 Netherlands (KY354230)
NE-1 consensus
ARV-1 Austria (OL803821)
ARV-1 Israel (MF114351)
0.002
New Zealand
United States
China
South Africa
Netherlands
B. ARV-2
NZ-4 consensus
NZ-5 consensus
NZ-9p consensus
NZ-2 consensus
ARV-2 Tonga (KY354234)
NZ-1 consensus
NZ-3 consensus
ARV-2 China (OM744320)
ARV-2 China (MZ821795)
ARV-2 China (OM744319)
ARV-2 China (MZ821796)
NE-6 consensus
NE-4 consensus
NE-3 consensus
NE-2 consensus
NE-5 consensus
NE-1 consensus
SA-2 consensus
SA-1 consensus
ARV-2 South Africa (KY354233)
SA-6p consensus
US-1p consensus
US-2p consensus
SA-5p consensus
0.002
C. VDV-3/-5
NZ-7 consensus
NZ-2 consensus
NZ-3 consensus
NZ-5 consensus
NZ-1 consensus
NZ-2 consensus
NZ-variant- (1*)
NZ-6 consensus
VDV-5 UK (MK795519)
NE-5 consensus
NE-8p consensus
CH-1p consensus
NE-1 consensus
NE-6 consensus
NE-7p consensus
NE-4 consensus
NE-8p consensus
SA-5p consensus
SA-6p consensus
SA-4 consensus
US-1p consensus
US-2p consensus
VDV-3 Israel (NC_040313)
CH-3p consensus
0.09
D. VDV-2
NE-3 Israel consensus
NE-4 Israel consensus
NE-6 Israel consensus
NE-2 Israel consensus
NE-8p Israel consensus
NE-7p Israel consensus
NE-5 Israel consensus
NE-1 Israel consensus
CH-3p Israel consensus
CH-1p Israel consensus
CH-2p Israel consensus
VDV-2 Israel reference (NC_040601)
NZ-7 Israel consensus
NZ-6 Israel consensus
NZ-3 Israel consensus
NZ-4 Israel consensus
NZ-2 Israel consensus
NZ-1 Israel consensus
NZ-9p Israel consensus
NZ-8p Israel consensus
NZ-5 Israel consensus
US-2p Israel consensus
US-1p Israel consensus
SA-1 Israel
SA-2 Israel
SA-4 Israel
SA-3 Israel
SA-6p Israel
SA-5p Israel
SA-3 UK
SA-4 UK
SA-5p UK
SA-6p UK
SA-1 UK
SA-2 UK
NE-3 UK consensus
NE-4 UK consensus
NE-6 UK consensus
NE-7p UK consensus
NE-5 UK consensus
NE-2 UK consensus
NE-8p UK consensus
NE-1 UK consensus
VDV-2 UK reference (MK795517)
CH-2p UK consensus
CH-3p UK consensus
CH-1p UK consensus
NZ-3 UK consensus
NZ-7 UK consensus
NZ-2 UK consensus
NZ-4 UK consensus
NZ-8p UK consensus
NZ-6 UK consensus
NZ-5 UK consensus
NZ-9p UK consensus
NZ-1 UK consensus
US-2p UK consensus
US-1p UK consensus
NZ-3 China consensus
NZ-2 China consensus
NZ-4 China consensus
NZ-7 China consensus
NZ-1 China consensus
NZ-6 China consensus
NZ-8p China consensus
NZ-9p China consensus
NZ-5 China consensus
NE-6 China consensus
VDV-2 China reference (MW590582)
CH-3p China consensus
CH-1p China consensus
CH-2p China consensus
NE-1 China consensus
NE-5 China consensus
NE-2 China consensus
NE-8p China consensus
NE-2 China consensus
NE-3 China consensus
NE-7p China consensus
SA-1 China
SA-2 China
SA-4 China
SA-6p China
SA 5p China
SA-3 China
US-2p China consensus
US-1p China consensus
0.09

**Figure S3.** Phylogenetic analysis of consensus sequences of *V. destructor- associated* viruses. Alignments were performed using consensus sequences generated from iterative mapping of V. destructor small RNA reads from each sample to **A)** Apis rhabdovirus 1 (ARV-1); **B)** Apis rhabdovirus 2 (ARV-2); **C)** Varroa destructor virus 3 & 5 (VDV-3/-5) and **D)** Varroa destructor virus 2 (VDV-2), using all three available reference strains from the UK (MK795517), Israel (NC_040601) & China (MW590582).
**A)** Whole genome consensus sequences of ARV-1 were aligned using MUSCLE and trimmed for gaps, leaving a final length of 12835 nt. The phylogenetic tree was generated using maximum likelihood in IQ-TREE [1], with the TIM+F+G4 model which had the optimal BIC score, as determined by ModelFinder [2]. **B)** ARV-2 was analysed as above, with final alignment length of 12718 nt and the TIM+F+I model. **C)** VDV-3 and VDV-5 were analysed as above with final alignment of 4082 nt and TIM2+F+G4 model. **D)** VDV-2 was analysed as above with final alignment of 8246 nt and GTR+F+I+G4 model. For A-D, branch supportsvwere estimated using Ultrafast bootstrap approximation (UFBoot; [3]) using 1000 replicates. Support values shown are SH-aLRT and UFBoot supports.

**1.** Nguyen, L.-T.; Schmidt, H.A.; von Haeseler, A.; Minh, B.Q. Iq-tree: A fast and effective stochastic algorithm for estimating maximum-likelihood phylogenies. Mol Biol Evol 2014, 32, 268-274.
**2.** Kalyaanamoorthy, S.; Minh, B.Q.; Wong, T.K.F.; von Haeseler, A.; Jermiin, L.S. Modelfinder: Fast model selection for accurate phylogenetic estimates. Nat Methods 2017, 14.
**3.** Hoang, D.T.; Chernomor, O.; von Haeseler, A.; Minh, B.Q.; Vinh, L.S. Ufboot2: Improving the ultrafast bootstrap approximation. Mol Biol Evol 2017, 35, 518-522.

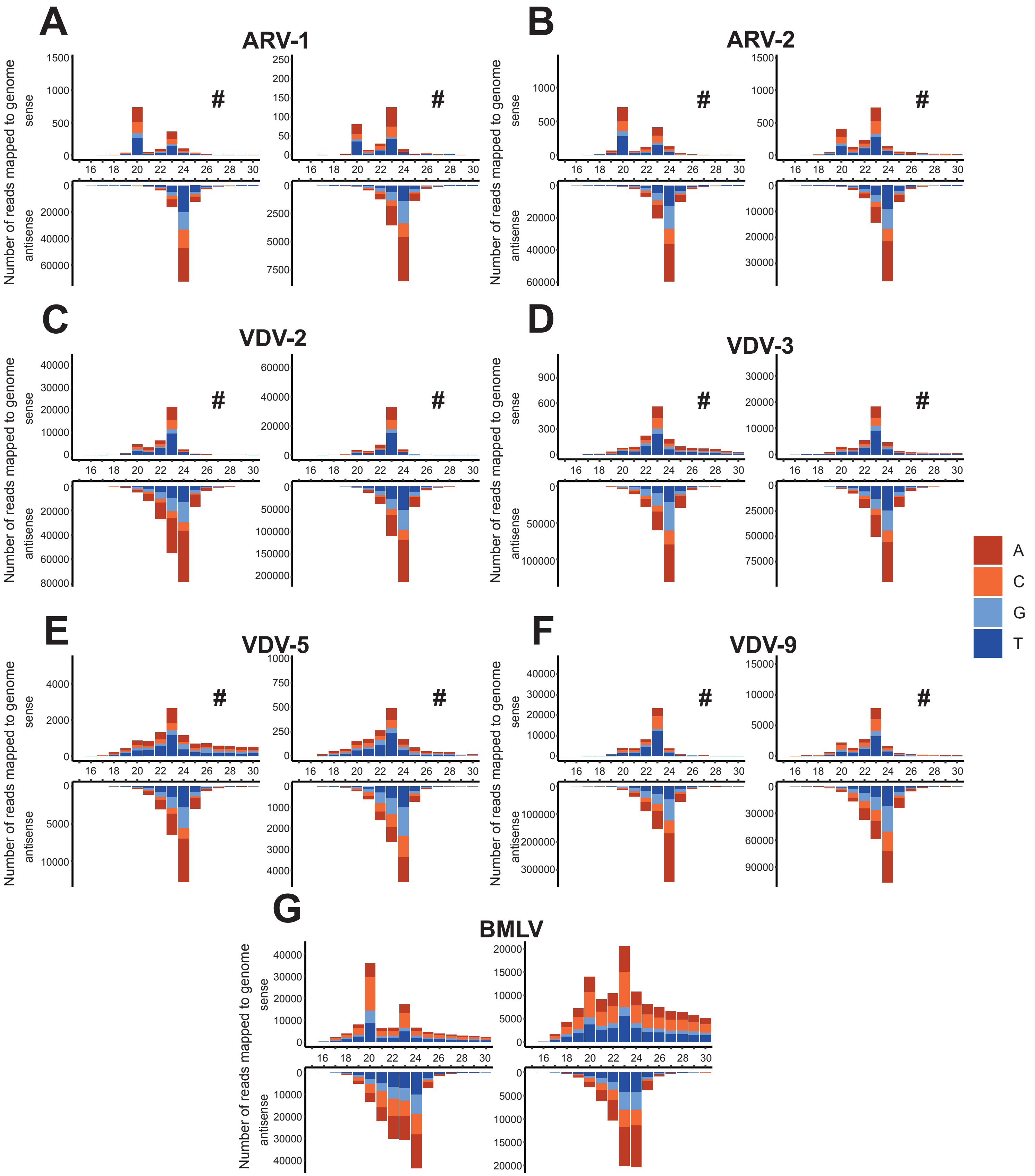


**Figure S4.** Additional samples showing representative size profiles of viral small RNA reads mapping to A) Apis rhabdovirus 1 (ARV-1: NE-5, SA-4); B) Apis rhabdovirus 2 (ARV-2: NE-2, SA-6p); C) Varroa destructor virus 2 (VDV-2: NE-8p, SA-5p); D) Varroa destructor virus 3 (VDV-3: CH-3, SA-5p); E) Varroa destructor virus 5 (VDV-5: NZ-8p, CH-1); F) Varroa destructor virus 9 (VDV-9: NZ-1, NE-1); G) Bee macula-like virus (BMLV: NE-6, NE-7p). # indicates sense Y-axis has been scaled to aid in visualising any size peaks.
